# Oncogenic Alterations, Race, and Survival in US Veterans with Metastatic Prostate Cancer Undergoing Somatic Tumor Next Generation Sequencing

**DOI:** 10.1101/2024.10.24.620071

**Authors:** Luca F. Valle, Jiannong Li, Heena Desai, Ryan Hausler, Candace Haroldsen, Monica Chatwal, Matthias Ojo, Michael J. Kelley, Timothy Rebbeck, Brent S. Rose, Matthew B. Rettig, Nicholas G. Nickols, Isla P. Garraway, Kosj Yamoah, Kara N. Maxwell

## Abstract

**Purpose:** National guidelines recommend next generation sequencing (NGS) of tumors in patients diagnosed with metastatic prostate cancer (mPCa) to identify potential actionable alterations. We sought to describe the spectrum and frequency of alterations in PCa-related genes and pathways, as well as associations with self-identified race/ethnicity, and overall survival in US Veterans.

**Patients and Methods:** This retrospective cohort study included Non-Hispanic Black (NHB) and Non-Hispanic white (NHW) Veterans with mPCa who obtained NGS through the Veterans Affairs National Precision Oncology Program. 45 genes in seven canonical or targetable mPCa pathways were evaluated in addition to TMB and MSI status. Multivariable logistic regression evaluated associations between race/ethnicity and genomic alteration frequencies. Cox proportional hazards models were used to determine associations between race/ethnicity, specific gene/pathway alteration, and overall survival.

**Results:** 5,015 Veterans with mPCa who had NGS conducted were included (1,784 NHB, 3,231 NHW). NHB Veterans were younger, had higher PSA at diagnosis, were less likely to report Agent Orange exposure, and resided in more deprived neighborhoods compared to NHW Veterans. Nine of the top ten most commonly altered genes were the same in NHB v NHW Veterans; however, the frequencies of alterations varied by race/ethnicity. NHB race/ethnicity was associated with higher odds of genomic alterations in *SPOP* (OR 1.7 [1.2-2.6]) as well as immunotherapy targets (OR 1.7 [1.1-2.7]) including MSI high status (OR 3.1 [1.1-9.4]).

Furthermore, NHB race/ethnicity was significantly associated with lower odds of genomic alterations in the AKT/PI3K pathway (OR 0.6 [0.4-0.7]), AR axis (OR 0.7 [0.5-0.9]), and tumor suppressor genes (OR 0.7 [0.5-0.8]). Cox proportional hazards modelling stratified by race/ethnicity demonstrated alterations in tumor suppressor genes including *TP53* were associated with shorter OS in both NHB (HR 1.54 [1.13-2.11] and NHW individuals (HR 1.52 [1.25-1.85]).

**Conclusion:** In the equal access VA healthcare setting, Veterans undergoing NGS for mPCa exhibited differences in alteration frequencies in both actionable and non-actionable pathways that may be associated with survival. This analysis affirms the utility of genomic testing for identifying candidates irrespective of race/ethnicity for precision oncology treatments, which could contribute to equitable outcomes in patients with mPCa.

## INTRODUCTION

The goal of precision oncology is to personalize prognostication and treatment for individual patients and to identify candidates for life-prolonging targeted therapies. Access to next generation sequencing (NGS) is essential to identify candidates for precision oncology approaches who are diagnosed with metastatic prostate cancer (mPCa), and national guidelines strongly recommend NGS in this heterogeneous patient population.

Racial disparities in PCa incidence and outcomes are well documented^1–4^. In the equal access Veterans Administration (VA) healthcare system, self-identified non-Hispanic Black (NHB) Veterans experience a higher incidence of localized and metastatic prostate cancer compared to non-Hispanic white (NHW) Veterans^5,6^. However, differences in survival remain unclear, especially in those with advanced disease^5,7^. NHB men are notably underrepresented in precision medicine cohorts^8^ including those that report genomic alteration frequencies^9,10^.

The National Precision Oncology Program within the Veterans Health Administration provisions NGS for Veterans with metastatic cancers^11^ and represents an unparalleled platform for assessing the landscape of alteration rates in mPCa across self-identified racial/ethnic groups in the diverse US Veteran population. In previous unadjusted analyses, differential rates of actionable alterations based on self-identified race/ethnicity could not be identified^12^.

Herein, we report an analysis of alteration rates in both individual genes as well as hallmark prostate cancer pathways and actionable gene groupings in NHB and NHW Veterans diagnosed with mPCa within the VA Healthcare System. Our goal was to evaluate the frequency of commonly reported genomic alterations, as well as associations with overall survival after adjusting for patient demographic, clinical, pathological, and social determinants of health indices.

## METHODS

### Patients

This is a retrospective cohort study of US Veterans who underwent somatic tumor testing for mPCa between January 2015-December 2023. DNA sequencing data from tissue or plasma were eligible for inclusion. Tissue biospecimens included prostate biopsies, radical prostatectomy specimens, and prostate cancer metastases. All specimens were sequenced with the Foundation Medicine platform (FoundationOne® CDx or FoundationOne®Liquid CDx).

When multiple specimens from the same patient were sequenced (n = 232), the first sequenced specimen was selected for analysis. Genomic data and prostate cancer clinical and pathological data from VA clinical sources were co-analyzed under VA Central IRB approved Study #1729212, with data elements imported from Study #1612627.

Race and ethnicity were self-reported by Veterans and race categories for the purpose of this analysis were defined as non-Hispanic Black/African American (NHB) and non-Hispanic White (NHW). For one patient, race was known but sample sequenced was not, so this patient was excluded from analyses involving tissue type.

### Genomic Analyses

Short variant, copy number alteration and rearrangement variant calls were provided by Foundation Medicine to the National Precision Oncology Program, annotated and classified by oncogenicity (**Supplemental Table 1**). Oncogenic alteration rates were determined for each gene and for groupings of genes into hallmark oncogenic pathways in prostate cancer (**Supplemental Table 2**). These included mismatch repair (MMR) deficiency genes, immunotherapy targets, DNA repair pathways, prostate cancer-specific PARP inhibitor (PARPi) targets, the AKT/PI3K pathway, AR signaling pathways, tumor suppressor pathways, and other known targetable pathways… Alterations affecting fewer than 11 patients were not reported to protect Veteran privacy.

### Statistical Analysis

Alteration frequencies were calculated by dividing the number of alterations identified for a specific gene or gene grouping by the total number of patients tested for that gene or gene grouping. Fisher’s exact testing was used to compare alteration frequencies based on Veteran self-identified race and specimen tested in our univariate analysis. p-values underwent false discovery rate correction for both the individual gene and gene grouping analysis, to account for multiple comparisons.

For genes where significant differences in alteration frequencies were identified between NHB and NHW Veterans, multivariable logistic regression was then carried out on those individual genes and their associated pathways to identify differential associations between race and alteration frequencies among all samples tested. Covariates that were adjusted for included sample type, age at diagnosis, age at sample collection, de novo metastatic status, castration resistance status, PSA level at diagnosis, Gleason grade, military exposures, pathological diagnosis, Charlson Comorbidity Index, Area Deprivation Index (ADI), smoking status, and marital status.

Adjusted overall survival estimates were evaluated using a Cox regression model, stratified by race and adjusted for the same above-mentioned covariates.

## RESULTS

Among a total of 6498 patients with a PCa diagnosis who received NGS testing through NPOP, 5559 patients self-identified as either NHB or NHW. After excluding 544 patients with non-metastatic disease using a natural language processing algorithm (previously described^13^), and one patient for lack of annotation of tissue type (primary vs metastatic site), sequencing data from 5015 Veterans with mPCa were available for analysis (**Supplemental Figure 2**).

Patient and disease characteristics from both the time of diagnosis are shown in **Table 1**. Patient and disease characteristics in relation to the time of NGS specimen collection are shown in **Supplemental Table 3**. 1,784 patients self-identified as NHB and 3,231 self-identified as NHW. NHB Veterans were significantly younger at the time of diagnoses, presented with higher PSAs at diagnosis, were less likely to have Agent Orange exposure, resided in state block groups with higher ADI, and were less likely to be deceased at the time of our analysis.

**Table 1:**
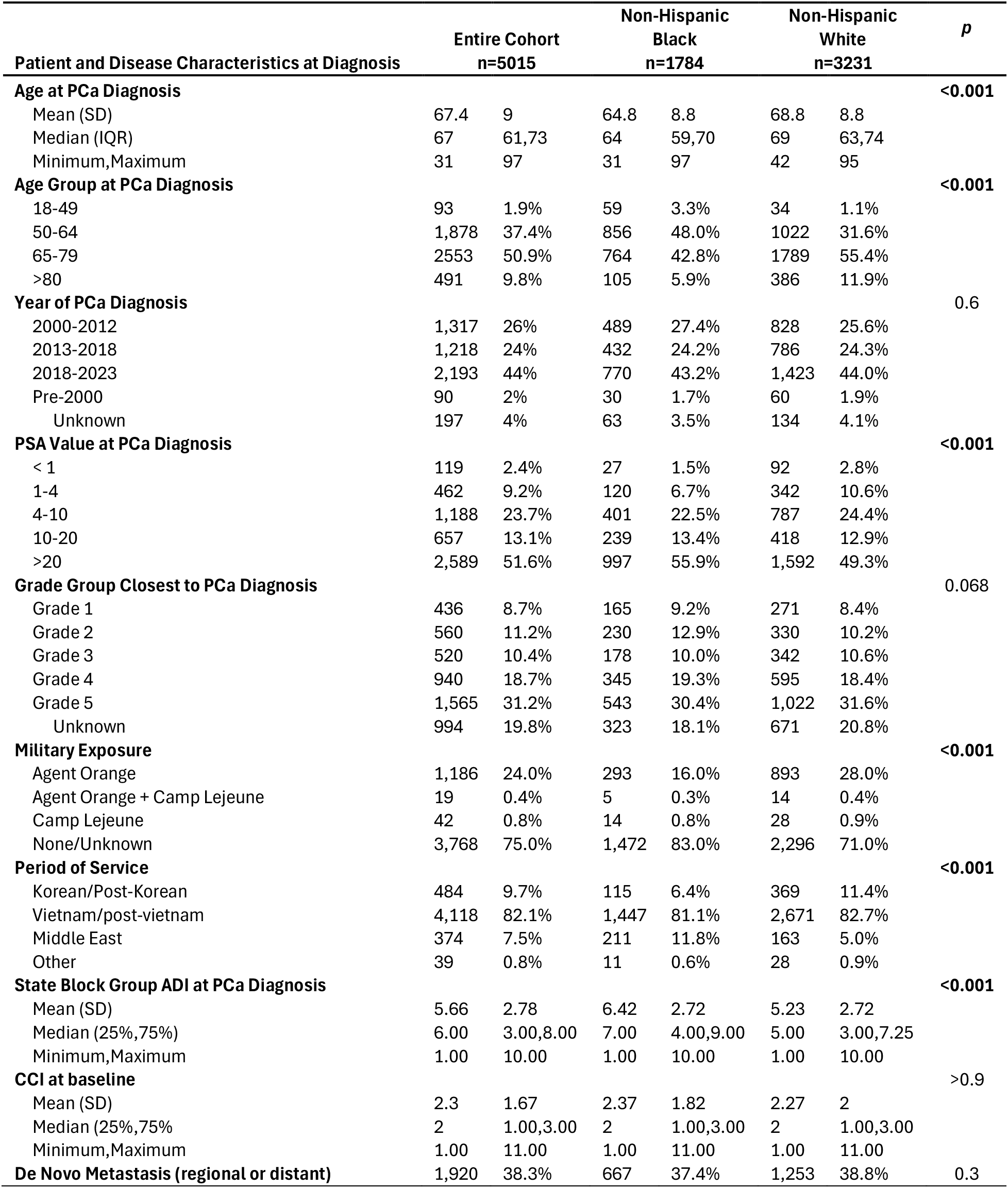
Patient and Disease Characteristics of Self-identified non-Hispanic Black (NHB) and non-Hispanic White (NHW) Patients with Metastatic Prostate Cancer and NGS Tumor Testing.

In an unadjusted analysis of all NGS analyte results combined, NHW Veterans were significantly more likely to harbor alterations in AKT/PI3K pathway genes (30% vs 20%, p<0.001), AR signaling axis genes (44% vs 35%, p<0.001), DNA repair genes (22% vs 17%, p<0.001), prostate cancer-specific PARPi targets (25% vs 22%, p=0.03), and tumor suppressor genes (52% vs 38%, p<0.001) (**Supplemental Figure 2A** and **Supplemental Table 4**). By contrast, NHB Veterans were significantly more likely to harbor alterations in immunotherapy targets (11% vs 7.2%, p<0.001), MMR genes (4.6% vs 3.0%, p<0.01), and were more frequently MSI high (4.7% vs 2.7%, p=0.029). There were no significant differences in TMB status or alterations in other targeted therapy pathways.

Upon interrogating alteration frequency by NGS analyte (plasma vs tissue) and tissue of origin (primary prostate vs metastatic site) without stratifying for self-identified race, we observed significant differences in alterations within the hallmark pathways, with metastases significantly more likely to harbor alterations in all pathways compared to primary tumor tissue (**Supplemental Table 5**). Plasma specimens were also more likely to demonstrate DNA repair alterations (30% vs 15%, p<0.001), prostate cancer-specific PARPi target alterations (31% vs 19%, p<0.001) and MSI high status (21% vs 2.3%, p<0.001) but significantly less likely to harbor alterations targetable by immunotherapy (6.0% vs 9.0%, p<0.001), AKT/PI3K pathway alterations (15% vs 29%, p<0.001), and AR signaling alterations (32% vs 42%, p<0.001) compared to primary tumors. Alterations in MMR, tumor suppressor genes, and other targetable pathways were no different between plasma and primary tissue. TMB status was low and no different across analytes (**Supplemental Figure 2B, Supplemental Table 5**).

Given the importance of race and NGS analyte on alteration frequencies, we carried out an aggregate comparison of alteration frequency stratified by both race and tissue type sampled, focusing on genomic alterations that were significantly different in the separate race and tissue-based analyses above. The most commonly altered genes were similar between NHB v NHW Veterans stratified by NGS analyte type, but the oncogenic alteration rates were significantly different between NHB v NHW Veterans for multiple genes (**Supplemental Figures 3-5; Supplemental Table 6**). AR axis alterations were significantly more likely to be identified in NHW samples derived from primary (46% vs 35%, p<0.001) and metastatic (56% vs 40%, p<0.001) tissue, however no differences were uncovered in plasma samples (33% vs 31%, p=0.61). The same trends of significance emerged for alterations in the AKT/PI3K Pathway (**Figure 1A, Supplemental Table 5=6**). By contrast, only in sequenced plasma samples was a significant increase in prostate cancer-specific PARPi target alterations (34% vs 27%, p=0.01) observed in NHW men. Alterations in DNA repair genes were more common in NHW Veterans compared to NHB Veterans when primary tissue (16% vs 11%, p=0.001) and plasma (33% vs 25%, p=0.005) were tested. Regarding alterations in tumor suppressor genes, NHW patients were significantly more likely to harbor alterations in this group of genes compared to NHB patients in all tissue types tested. Alterations in immunotherapy pathways (13% vs 6.8%, p<0.001), dMMR (5.3% vs 2.4%, p<0.001), and other known targetable pathways (10% vs 5.9%, p<0.001), were all significantly more likely in NHB patients, but only when the primary tumor was tested (**Figure 1A, Supplemental Table 6**).

**Figure 1:**
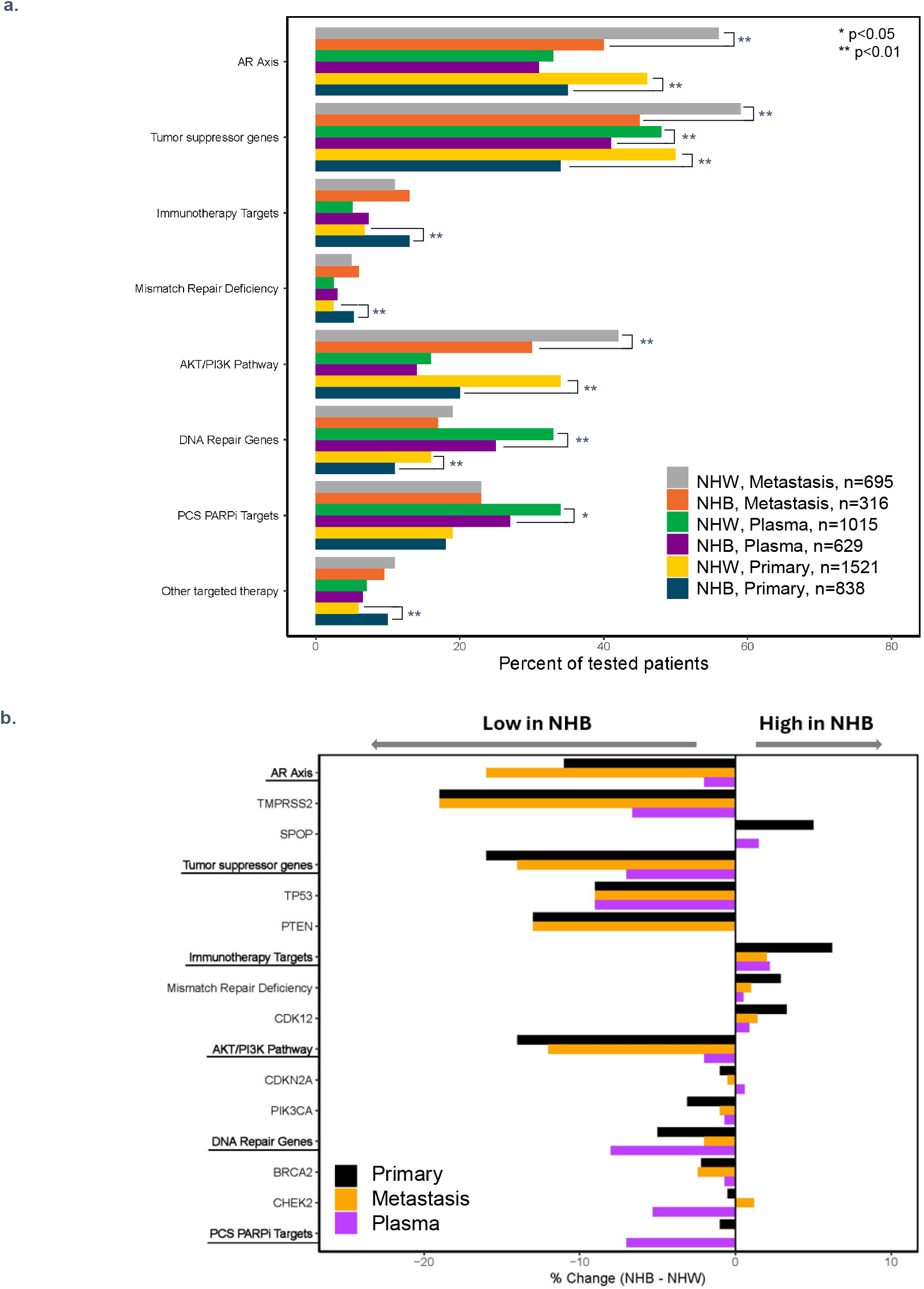
Association of genomics, race and survival in metastatic prostate cancer patients. a. Rates of oncogenic alterations in genes from cancer-related pathways in three different tissues (primary tumors, metastatic lesions and plaslmasamples), separated by patient self-identified race in individuals with metastatic prostate cancer. Differences in rates of oncogenic alterations between NHB and NHW patients were compared by Fisher’s exact test for each of the three tissue types. b.Percent change in oncogenic alteration rates in pathways their constituent genes within those pathways in NHB compared to NHW patients in three tissue types. NHB = non-Hispanic Black; NHW = non-Hispanic White, AR = androgen receptor, PCS = prostate cancer specific, PARPi= PARP inhibitor.

When considering alterations in the individual genetic components of the previously described oncogenic pathways that were significantly different by race, it becomes clear that AR signaling differences are likely driven by alteration rates in *TMPRSS2* rather than *SPOP* (**Figure 1B**). *TP53* and *PTEN* alterations appear to contribute similarly to the lower tumor suppressor alteration frequency in NHB Veterans, while dMMR and *CDK12* alterations appear to contribute similarly to the increase in immunotherapy targets in NHB individuals (**Figure 1B**).

Since analyte and race can both impact alteration frequencies, multivariate analysis was pursued to better understand the association between race and alteration frequency. After multivariable adjustment for patient and tumor-related characteristics including NGS analyte, clinicopathologic features, and social determinants of health (SDOH) covariates, alteration frequencies in NHB Veterans were found to be significantly lower in AR axis genes (OR 0.7, p<0.01), tumor suppressor genes (OR 0.7, p<0.001), DNA repair genes (OR 0.7, p<0.01), and AKT/PI3K pathway genes (OR 0.6, p<0.001), but significantly elevated in targets for immunotherapy (OR 1.7, p=0.02) (**Figure 2, Supplemental Table 7**).

**Figure 2:**
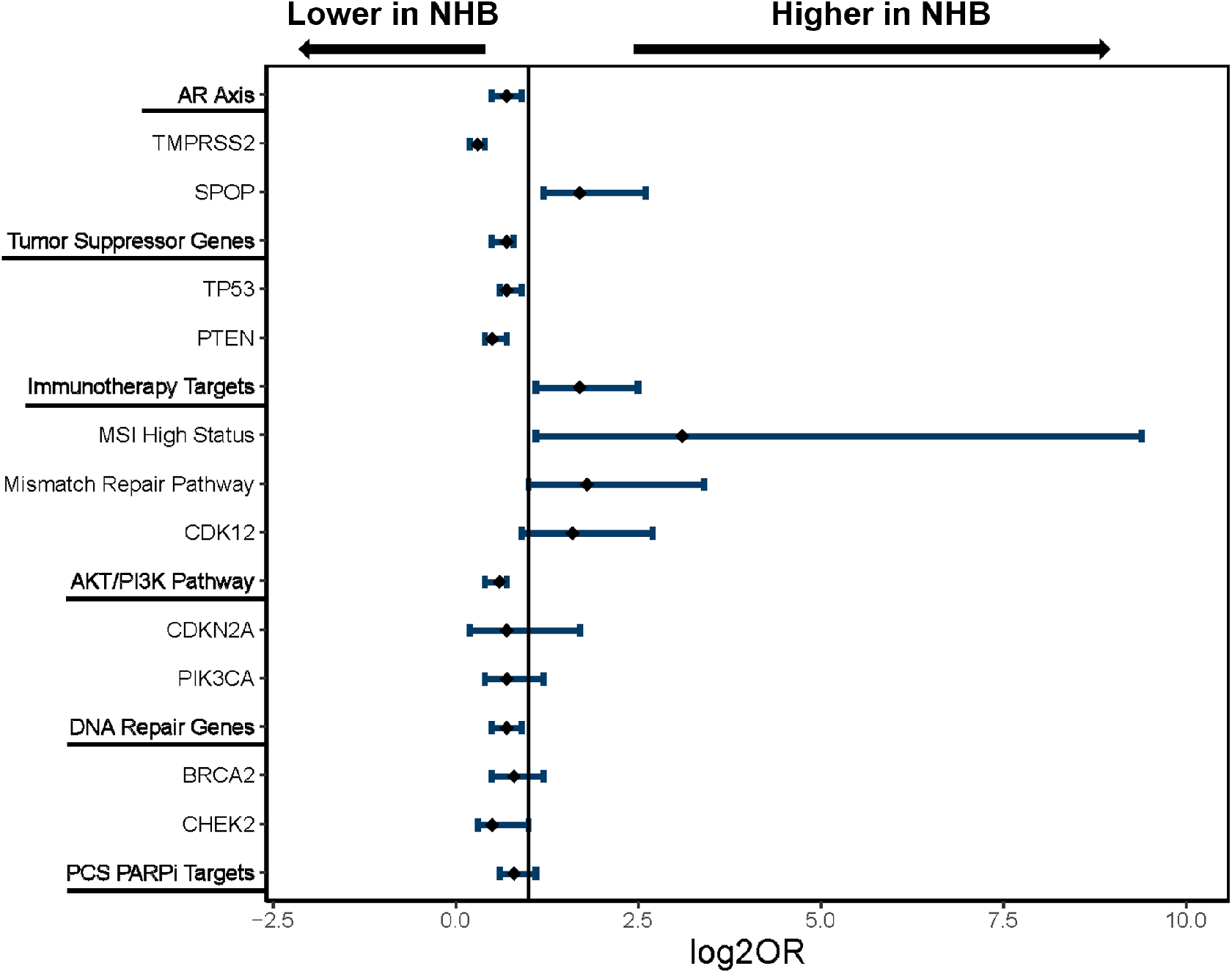
Association of race and alteration frequency in metastatic prostate cancer patients. Multivariate logistic regression analysis to evaluate the association of oncogenic alterations in pathways and genes in those pathways with self-identified race, controlling for multiple clinical factors. Pathways and genes were tested if statistically significantly difference in rates between Blacks and Whites in univariate analyses. NHB = non-Hispanic Black; NHW = non-Hispanic White, AR = androgen receptor, MSI = microsatellite instability, PCS = prostate cancer specific, PARPi = PARP inhibitor, OR = Odds Ratio.

Survival has been previously reported to be similar in NHB vs NHW Veterans who receive their care within the Veterans Health Administration^5,14^. In our adjusted Cox model for overall survival, tumor suppressor alterations (driven by *TP53* alterations), immunotherapy targets (driven by mismatch repair alterations and MSI high status), and AR axis alterations all increased the hazard of death in NHW Veterans, whereas tumor suppressor alterations (again driven by *TP53* alterations) and *CDK12* alterations all increased the hazard of death in NHB Veterans (**Figure 3, Supplemental Table 8**).

**Figure 3:**
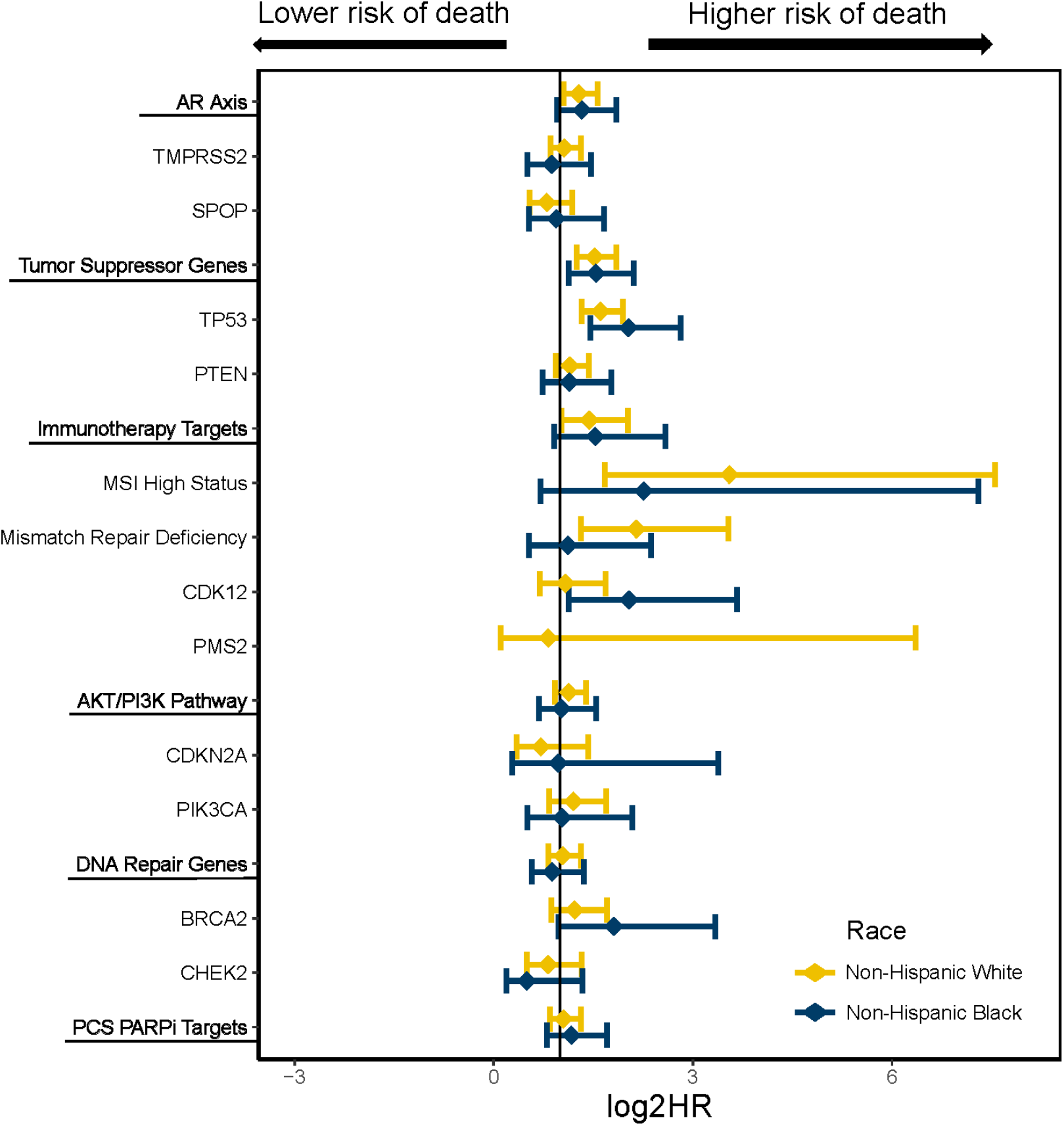
Association of genomics, race and survival in metastatic prostate cancer patients. Cox proportional hazards modeling to test the association of oncogenic alterations in pathways and genes in those pathways with overall survival, controlling for clinical factors and stratified by race, NHB = non-Hispanic Black; NHW = non-Hispanic White, AR = androgen receptor, MSI = microsatellite instability, PCS = prostate cancer specific, PARPi = PARP inhibitor, HR = Hazard Ratio.

## DISCUSSION

This report describes alteration frequencies in key PCa pathways and genes, including those that are known to be targetable with precision oncology interventions in a large cohort of NHB and NHW Veterans with mPCa. Absolute and proportional (36%) representation of NHB individuals on this study was markedly higher than previous reports^9,10,15,16^. The observed increased frequency of potential immunotherapy targets in NHB individuals is of interest because of potential actionability in this population and underscores the continued need for equitable application of precision medicine efforts in mPCa, as this could be one of many factors that can mitigate disparities in PCa outcome.

Associations between responsiveness to immunotherapy and patient race have been previously reported from real-world PROCEED registry data^17^, where OS was significantly higher in Black men with mCRPC who were treated with the first FDA approved cellular immunotherapy, sipuleucel-T. Examples of associations between MSI high status and complete response to sipuleucel-T have been reported in the literature, and MSI high status is also a marker for responsiveness to PD-1 axis inhibition^18^, so it is possible that the higher frequency of alterations in immunotherapy targets in NHB men, driven by higher MSI, may be contributing to enhanced immunogenicity of tumors in this population. Consequently, this heightened immunogenicity could potentially lead to more favorable responses to immunotherapy. Of note, TMB, a known predictor of response to checkpoint inhibition, was not significantly different between NHW and NHB Veterans. Our finding that NHW men with PCa have a higher frequency of *TP53* alterations aligns with the hypothesis proposed by Halabi and colleagues suggesting an interaction between an intact *TP53* pathway and the efficacy of docetaxel as a plausible explanation for the improved outcomes observed in Black men treated with the wild-type p53 dependent drug, docetaxel^19^.

In addition to our findings related to immunotherapy targets, NHW individuals were more likely to harbor alterations in AR signaling than NHB individuals. This may contribute to decreased responsiveness to androgen receptor signaling inhibitors (ARSI) in NHW compared to NHB patients – an observation which has been reported in individuals with castrate resistant disease who received abiraterone as first line therapy^20^. This work also raises the possibility that while canonical AR activity may be lower in NHB patients, the activity of non-canonical AR pathways may be increased in NHB individuals, a possibility that requires further research.

Our report also underscores the fact that the source of tumor DNA (ie primary vs metastatic vs plasma) submitted for NGS testing can influence the frequency of specific alterations and confound comparisons between race, and thus remains an important control for analyses of this nature. Accordingly, sequencing results used by clinicians should always be considered in the context of the analyte tested. For example, our findings highlight that plasma evaluating cfDNA may not be as sensitive as tumor tissue for the identification of actionable immunotherapy targets; indeed, <10% of the plasma samples tested in this study reported microsatellite status. Plasma may also over-report frequencies of alterations in actionable DNA damage repair (DDR) genes (e.g. of *ATM* or *CHEK2*), which may be derived from clonal hematopoiesis especially in elderly patients^21^ such as Veterans with metastatic prostate cancer. Conversely, alterations in *CDKN2A, ERBB2, PTEN, RB1*, and *TMPRSS2* were found at lower frequencies in plasma, likely due to challenges identifying copy number variants and rearrangements in cfDNA versus tissue.

It has also been previously reported that many actionable PCa alterations, including DDR alterations, are truncal in nature^22^, and that their frequencies are therefore similar in both primary and metastatic tissue. Our work is largely consistent with this observation, and while metastases were significantly more likely to yield actionable alterations in DNA repair genes than primary tissue, this difference was small (24% vs 20%).

Prior unadjusted analyses have suggested that actionable alterations appear at similar rates between NHB and NHW Veterans whereas this larger adjusted analysis confirms DDR alteration frequencies are lower in NHB individuals, consistent with similar findings in the germline^23^. This adjusted analysis also suggests that immunotherapy targets are more frequent in NHB men. While this was certainly the case for MSI high status, the picture is less clear for *CDK12* alterations, which are known to be associated with aggressive PCa phenotypes. Our previous work revealed similar *CDK12* alteration rates between NHB and NHW Veterans ^12^, and while there was a higher frequency of these alterations in NHB Veterans in our univariate analysis, this difference disappeared after multivariable adjustment. Nevertheless, *CDK12* may still remain a consequential target for NHB men with mPCa, and we echo others^24,25^ in highlighting the fact that therapeutic targeting of CDK12 may serve as an avenue for improving outcomes for NHB patients.

A limitation of our analysis is the lack of matched germline data for these patients, which complicates interpretation of plasma results. Additionally, by analyzing the first specimen sent for NGS testing, there is a possibility of false negative reporting in the minority of patients who had multiple samples sent for NGS testing over the course of their disease, although the low frequency of these false negative cases would not significantly change our results or their interpretation.

Overall, this work emphasizes the notion that personalized precision medicine-driven approaches will be essential to maximally leverage our understanding of a patient’s specific tumor biology in order to improve oncologic outcomes. Even in scenarios where non-targetable alterations are identified (such as the case of tumor suppressor alterations including *TP53*), knowledge of the deleterious implications of these alterations on survival (consistent with previously reported work from Velez and colleagues^26^) can have wide implications for the management of both NHB and NHW men with mPCa and result in changes in treatment.

Specifically, given the observation that high-risk alterations (including *TP53, RB1*, and *PTEN*) can promote lineage plasticity, resulting in radiographic progression in the absence of PSA progression ^27,28^, the presence of these pathogenic alterations may have implications for the optimal cadence and modality of disease monitoring. Knowledge of these alterations may also assist in patient selection for metastasis-directed therapy (MDT) with stereotactic body radiotherapy (SBRT) in the oligometastatic setting, given the results of a recent pooled analysis that demonstrated a high-risk mutational signature consisting of alterations in BRCA1/2, ATM, TP53, or Rb1 were independently predictive of relative benefit and prognostic^29^ for response to MDT. And finally, this work implicitly advocates for the intentional design of precision medicine studies that do not exclude groups of patients that have been historically under-represented in clinical trials.

Precision oncology enables the individualization of treatment decisions without having to rely on imprecise characteristics such as self-identified race. We demonstrate that alteration frequencies in several hallmark oncogenic pathways in mPCa vary by race and that the association between specific alterations (e.g. *CDK12*, AR axis, dMMR) and survival, when stratified by race, is also variable. Thus, we did not identify any genetic alterations or biomarkers that should not be tested in prostate cancer on the basis of patient self-identified race. While individuals may exhibit different biological aggressiveness and associated outcomes from PCa treatment, precision-based testing and treatment approaches remain critical for personalizing care, optimizing outcomes, informing the design of equitable clinical trials, and narrowing disparities in outcomes for PCa.

## Author Contributions

Drs Garraway, Yamoah and Maxwell had full access to all of the data in the study and take responsibility for the integrity of the data and the accuracy of the data analysis. Drs Garraway, Yamoah and Maxwell contributed equally to this work and share last authorship/corresponding authorship.

***Concept and design:*** LFV, MBR, NGN, IPG, KY, KNM

***Acquisition, analysis, or interpretation of data:*** LFV, JL, HD, RH, CH, MC, MO, MJK, IPG, KNM

***Drafting of the manuscript:*** LFV, MBR, NGN, IPG, KY, KNM

***Critical review of the manuscript for important intellectual content:*** All authors

***Statistical analysis:*** JL, HD, RH

***Obtained funding:*** TR, BSR, MBR, NGN, IPG, KY, KNM

***Administrative, technical, or material support:*** N/A

***Supervision:*** TR, BSR, MBR, NGN, IPG, KY, KNM

## Disclaimer

The ideas and opinions expressed herein are those of the authors and do not necessarily reflect the opinions of the Veterans Affairs or their contractors and subcontractors.

