## Supplemental Figures for "Oncogenic Alterations, Race, and Survival in US Veterans with Metastatic Prostate Cancer Undergoing Somatic Tumor Next Generation Sequencing"

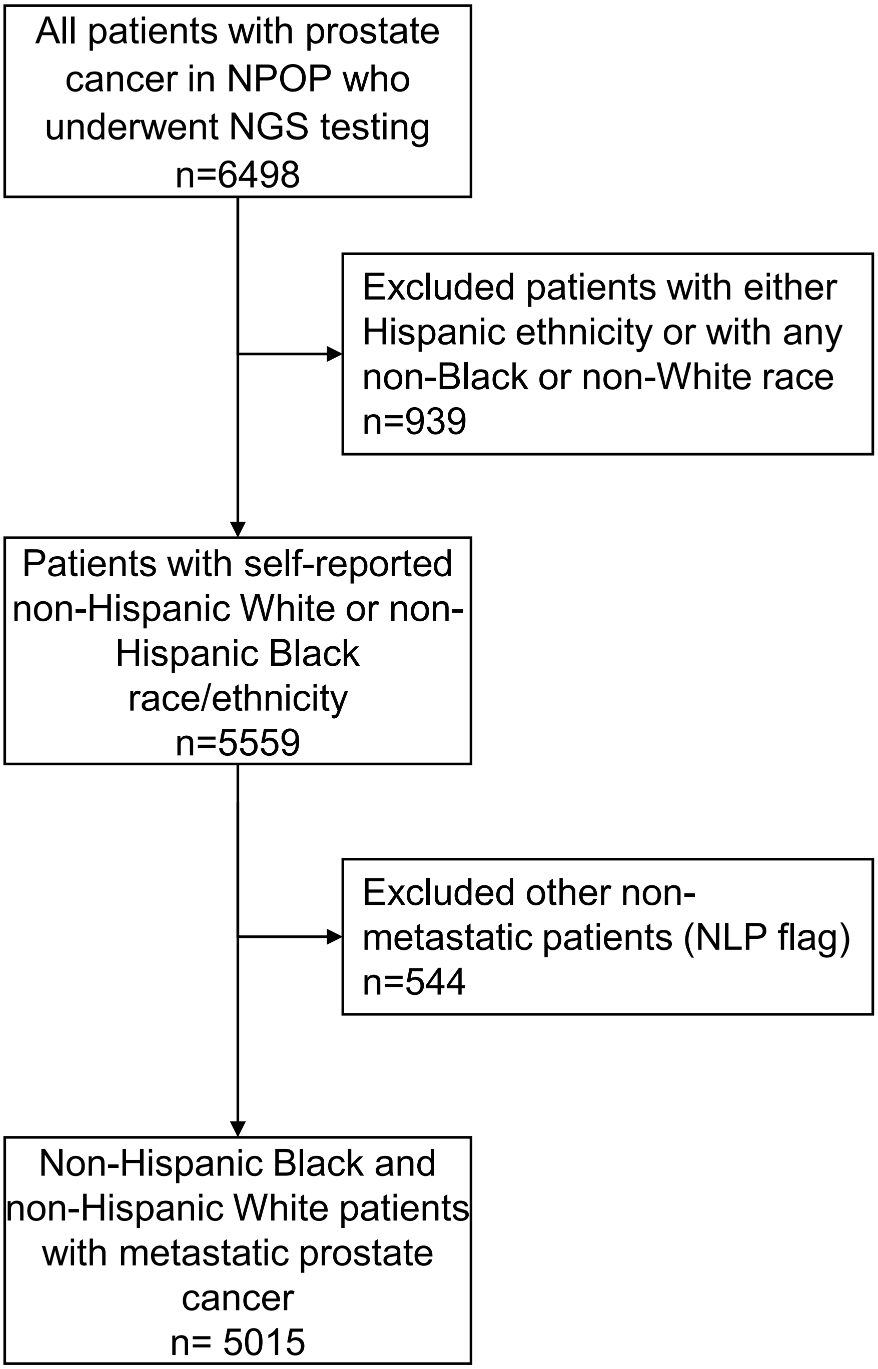


**Supplementary Figure 1:** Consort Diagram demonstrating patients included and excluded for analysis. NLP = natural language processing.

a.

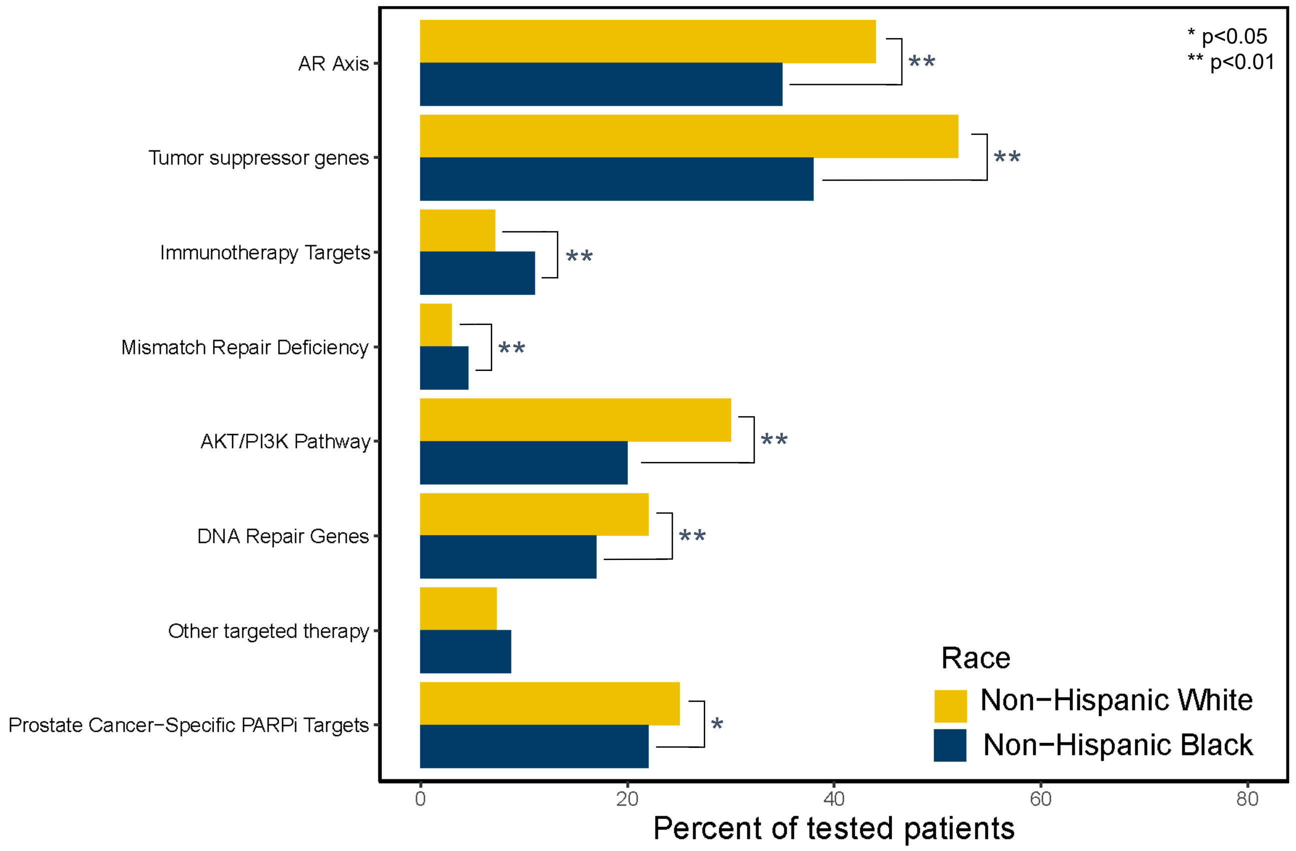


b.

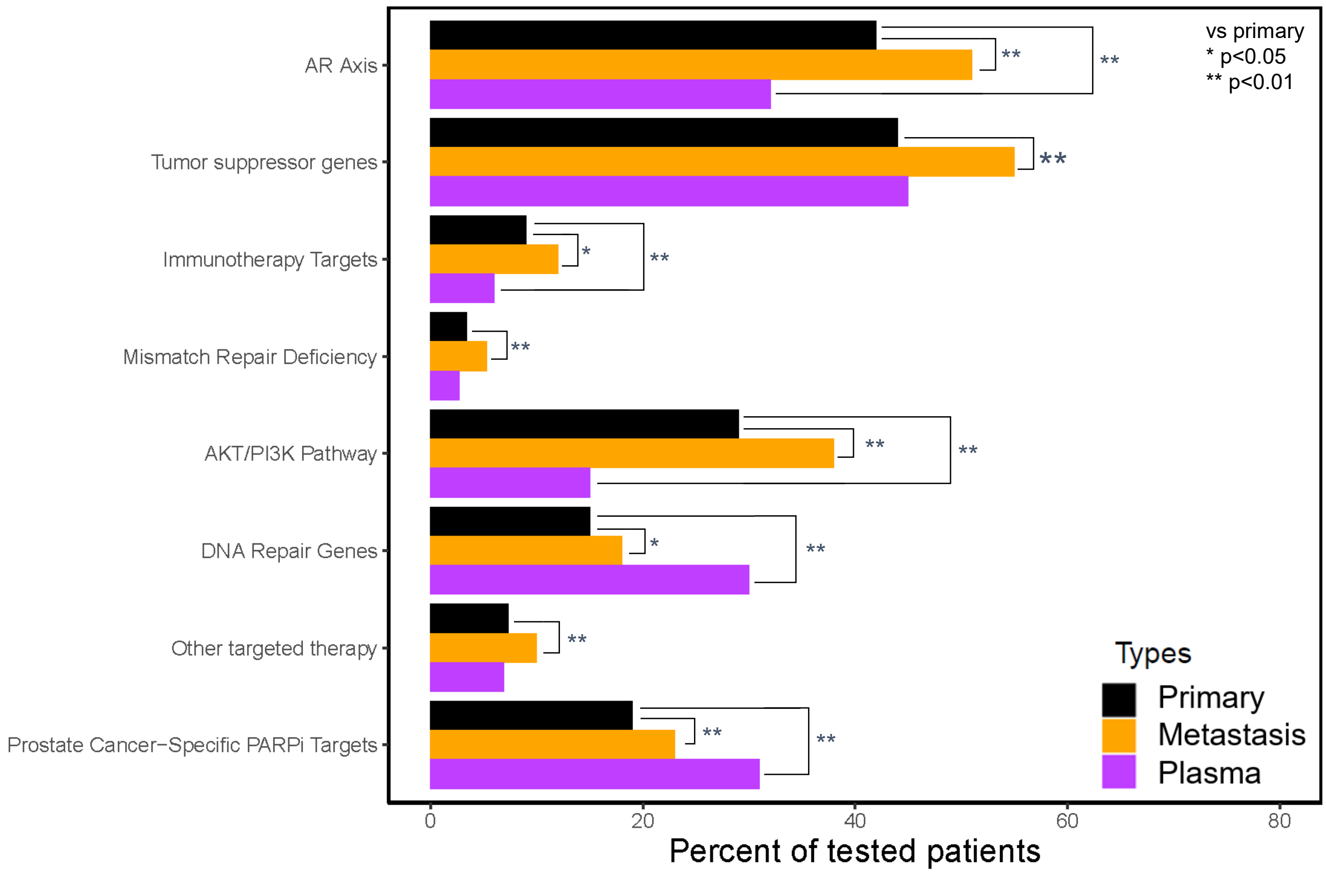


**Supplementary Figure 2: Differences in oncogenic pathways for individuals with metastatic prostate cancer organized by** A. patient self-identified race and by B. tissue analyzed for NGS testing. Genes involved in these pathways are listed in the Supplemental tables. NHB = non-Hispanic Black; NHW = non-Hispanic White, AR = androgen receptor, PCS = prostate cancer specific, PARPi = PARP inhibitor.

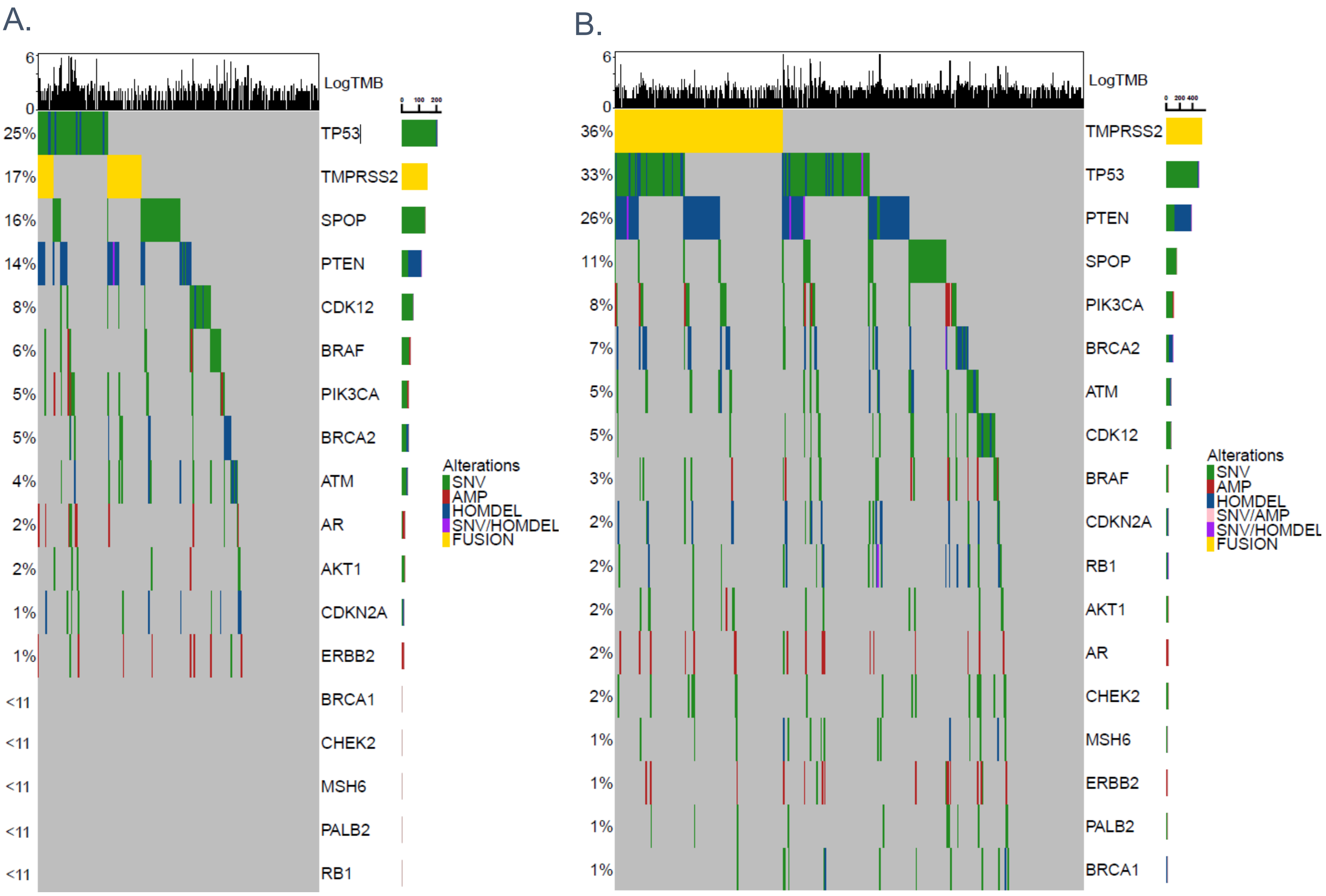


**Supplementary Figure 3:** Heatmaps denoting genomic alterations from primary tissue in A) NHB (n=838) and B) NHW (n=1521) Veterans.

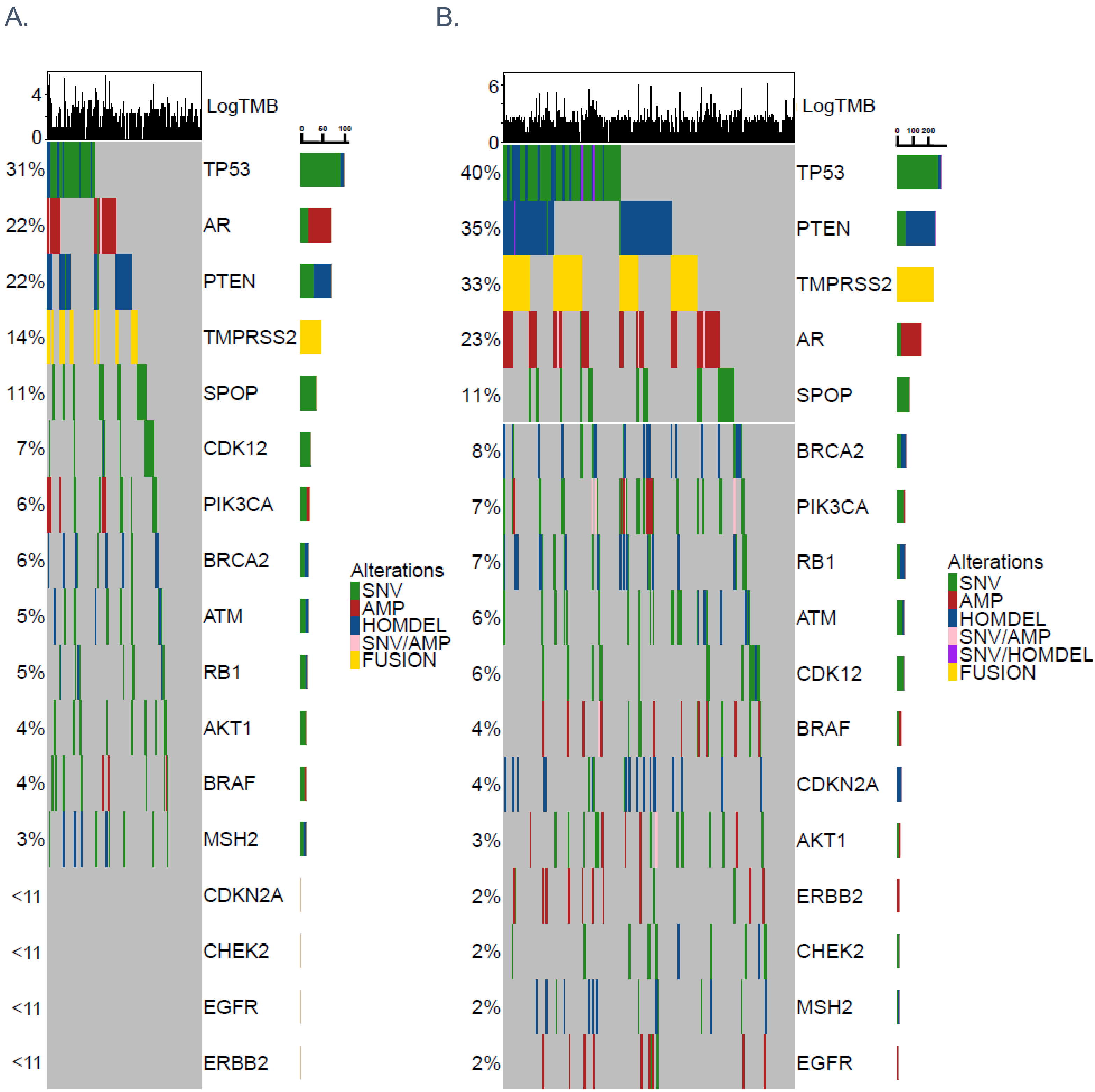


**Supplementary Figure 4:** Heatmaps denoting genomic alterations from metastatic tissue in A) NHB (n=316) and B) NHW (n=695) Veterans.

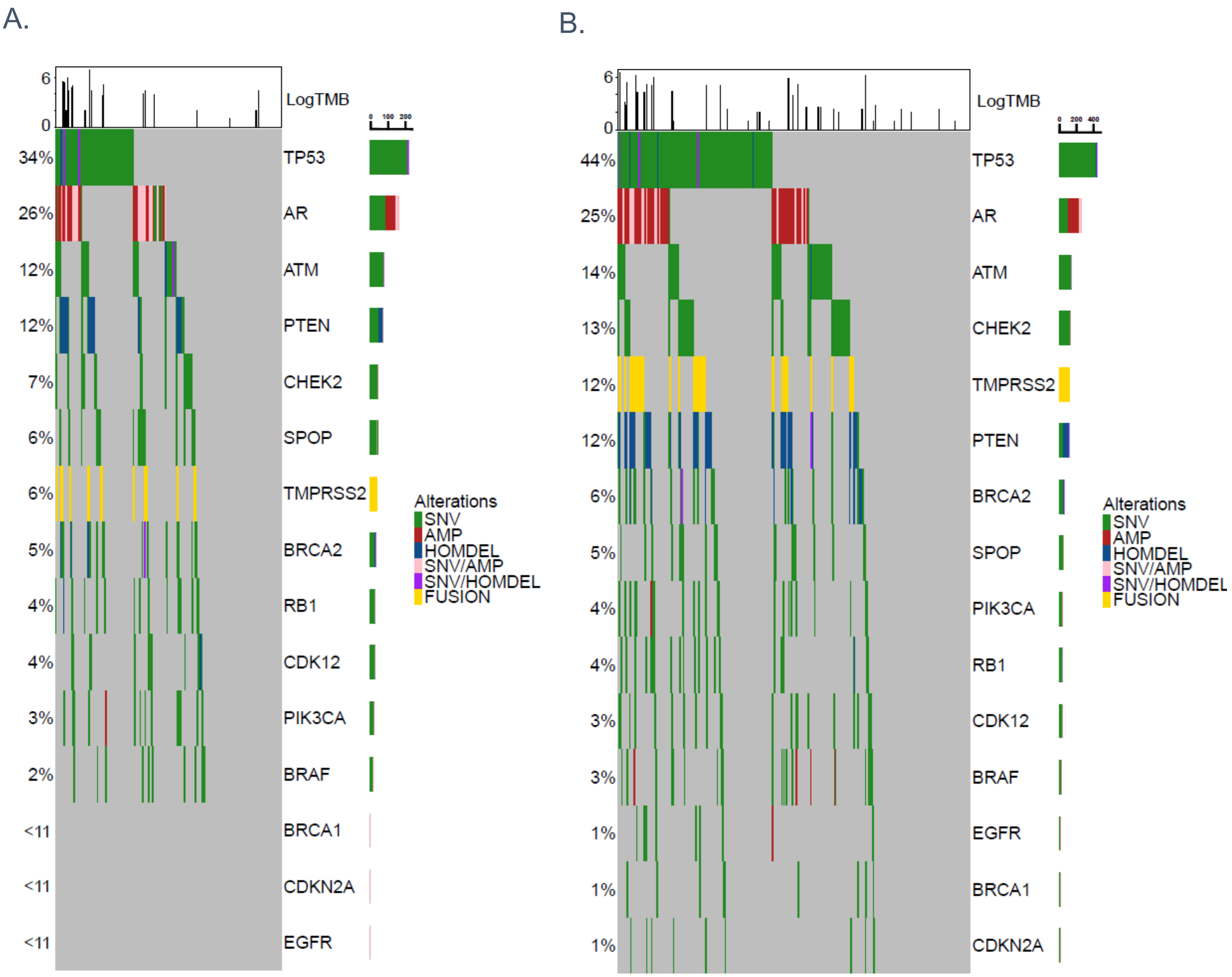


**Supplementary Figure 5:** Heatmaps denoting genomic alterations from liquid biopsy in A) NHB (n=629) and B) NHW (n=1015) Veterans.
